# Gene model for the ortholog of *Glyp* in *Drosophila ananassae*

**DOI:** 10.1101/2025.07.18.665411

**Authors:** Gabriella N. Bicanovsky, Evan N. Bennett, Zachary Lill, Rebecca M. McSweeney, Geoffrey D. Findlay, Amy T. Hark, Chinmay P. Rele, Shallee T. Page

## Abstract

We present a gene model for the ortholog of *Glycogen phosphorylase* (*Glyp*) in the *D. ananassae* May 2011 (Agencourt dana_caf1/DanaCAF1) genome assembly of *Drosophila ananassae* (GenBank Accession: GCA_000005115.1). This ortholog was characterized as part of a developing dataset to study the evolution of the insulin/insulin-like growth factor signaling pathway (IIS) across the genus *Drosophila* using the Genomics Education Partnership gene annotation protocol for Course-based Undergraduate Research Experiences.

## Introduction

This article reports a predicted gene model generated by undergraduates using a structured gene model annotation protocol defined by the Genomics Education Partnership (GEP; thegep.org) for Course-based Undergraduate Research Experiences (CUREs).

“In this GEP CURE protocol, students use web-based tools to manually annotate genes in non-model *Drosophila* species based on orthology to genes in the well-annotated model organism fruitfly *Drosophila melanogaster*. The GEP uses web-based tools to allow undergraduates to participate in course-based research by generating manual annotations of genes in non-model species (Rele et al., 2023). Computational-based gene predictions in any organism are often improved by careful manual annotation and curation, allowing for more accurate analyses of gene and genome evolution (Mudge & Harrow, 2016; Tello-Ruiz et al., 2019). These models of orthologous genes across species then provide a reliable basis for further evolutionary genomic analyses when made available to the scientific community in public databases.” (Myers et al., 2024).

The *D*.*ananassae Glyp* ortholog described here was characterized “…as part of a developing dataset to study the evolution of the insulin/insulin-like growth factor signaling pathway (IIS) across the genus *Drosophila*. The IIS pathway is a highly conserved signaling pathway in animals and is central to mediating organismal responses to nutrients (Grewal, 2009; Hietakangas & Cohen, 2009).” (Myers et al., 2024). “Glycogen phosphorylase (*Glyp*; also known as GP, Glp1, DGPH, FBgn0004507), part of the insulin signaling pathway, has a high degree of amino acid similarity to mammalian enzymes (Tick et al., 1999), and contains repeats that are similar to repeats in several transmembrane proteins including Notch in *D. melanogaster* and *Lin-28* in *Caenorhabditis elegans* (LaMarco et al., 1991). GLYP plays a major role in glycolytic activity in skeletal muscle in association with glycogen metabolism and body homeostasis (Yamada et al., 2019). The importance of *Glyp* is evident in that null homozygotes are lethal as full knockouts at the larval stage (Eanes et al., 2006; Yamada et al., 2019). The [Glyp gene product] has also shown to affect development, adult fitness, and aging in *Drosophila* (Bai et al., 2013; Yamada et al., 2019).” (Laskowski L.F. et al., 2025).

“*D. ananassae* (NCBI:txid7217) is part of the *melanogaster* species group within the subgenus *Sophophora* of the genus *Drosophila* (Sturtevant, 1939). First described by Doeschall (Doleschall, C.L., 1858), *D. ananassae* is circumtropical (Markow & O ‘Grady, 2006), and is often associated with human settlement (Singh, 2010). It has been extensively studied as a model for its cytogenetic and genetic characteristics, and in experimental evolution (Kent et al., 2002).” (Lawson et al., 2024).

### Synteny

The reference gene, *Glyp*, occurs on chromosome 2L in *D. melanogaster* and is flanked upstream by *Odorant receptor 22c* (*Or22c*) and *CG12674*, nested by *defective proboscis extension response 3* (*dpr3*), and downstream by *tho2* and *CG11723*. The *tblastn* search of *D. melanogaster* Glyp-PA (query) against the *D. ananassae* (GenBank Accession: GCA_000005115.1) Genome Assembly database placed the putative ortholog of *Glyp* within scaffold_12916 (European Nucleotide Archive: CH902620.1) at locus LOC6497423 (XP_001962117.1)— with an E-value of 0.0 and a percent identity of 94.58%. Furthermore, the putative ortholog is flanked upstream by LOC6497421 (XP_001962120.1), LOC26513848 (XP_014762266.1) and LOC6498121 (XP_044571907.1), which correspond to *Or22c, CG12674* and *dpr3* in *D. melanogaster* (E-value: 0.0, 5e-109 and 0.0; identity: 85.21%, 50.31% and 72.63%, respectively, as determined by *blastp*; Figure 1A; Altschul et al., 1990). The putative ortholog of *Glyp* is flanked downstream by LOC6498120 (XP_032307372.1) and LOC6498119 (XP_001962115.2), which correspond to *tho2* and *CG11723* in *D. melanogaster* (E-value: 0.0 and 5e-129; identity: 86.37% and 53.97%, respectively, as determined by *blastp*). The putative ortholog assignment for *Glyp* in *D. ananassae* is supported by the following evidence: The genes surrounding the *Glyp* ortholog are orthologous to the genes at the same locus in *D. melanogaster* and local synteny is completely conserved, supported by results generated from *tblastn* and *blastp*, so we conclude that LOC6497423 is the correct ortholog of *Glyp* in *D. ananassae* (Figure 1A).

**Figure 1:**
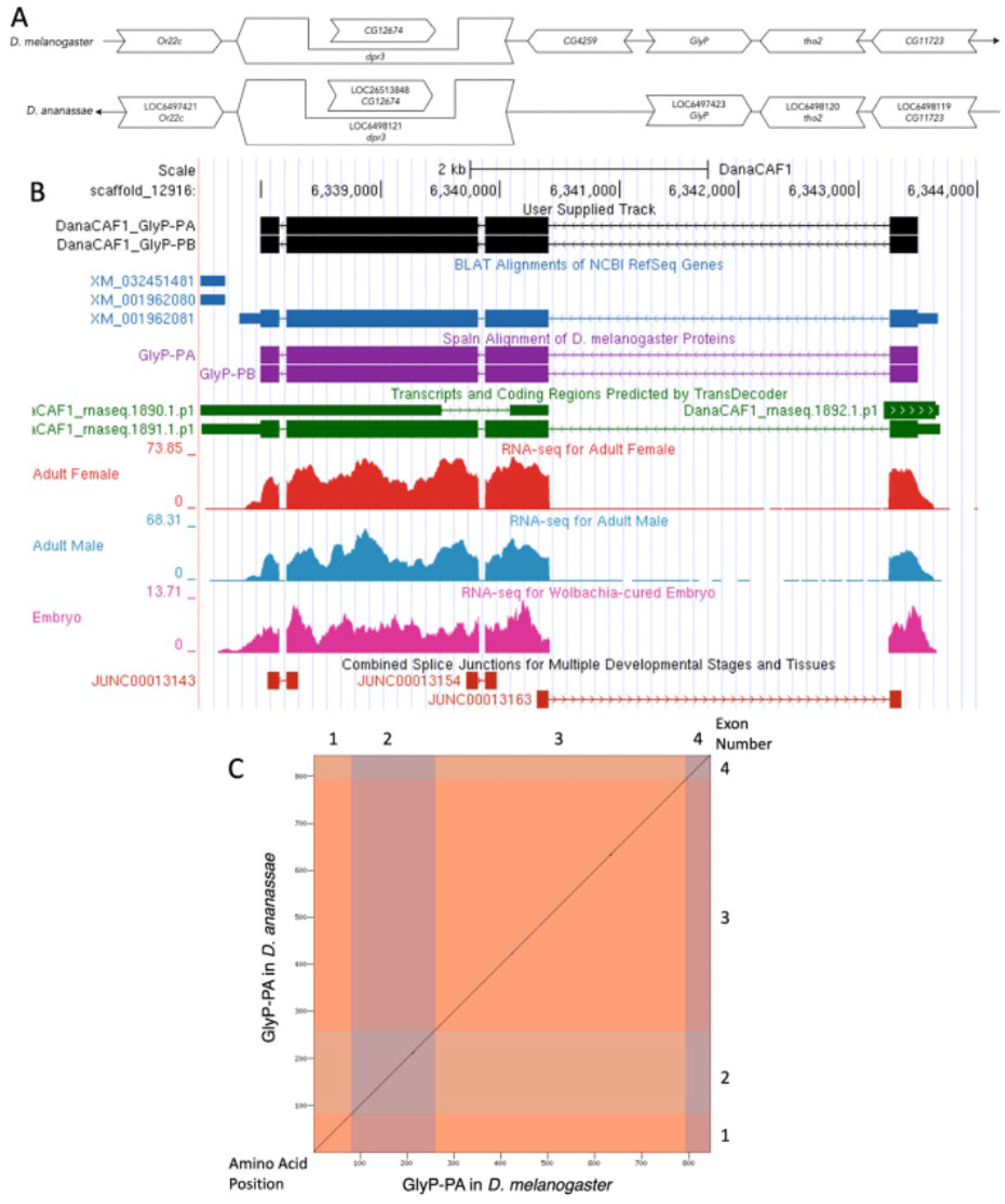
*Glyp* gene model comparison between *D. melanogaster* and *D. ananassae*: **(A) Synteny of the genomic neighborhood of *Glyp* in *D. melanogaster* and *D. ananassae***. The thin terminal arrow pointing to the right indicates that *Glyp* is on the (+) strand in *D. melanogaster*; the arrow pointing to the left indicates that *Glyp* is on the (–) strand in *D. ananassae*. Notations within the large white arrows in *D. ananassae* indicate the locus ID and the orthology to the corresponding gene in *D. melanogaster*. The wide gene arrows pointing in the same direction as *Glyp* are on the same strand relative to the thin underlying arrows, while wide gene arrows pointing in the opposite direction of *Glyp* are on the opposite strand. The embedded arrow indicates that the *Dpr-3* gene contains a nested gene (CG12674). The gene names given in the *D. ananassae* gene arrows indicate the orthologous gene in *D. melanogaster*, while the locus identifiers are specific to *D. ananassae*. **(B) The custom gene model in UCSC Track Hub** (Raney et al., 2014). The gene model in *D. ananassae* is depicted in black with exons depicted with wide boxes, introns with narrow lines, and arrows indicating direction of transcription. Evidence tracks below include BLAT alignments of NCBI Ref-Seq Genes (blue, alignment of Ref-Seq genes for *D. ananassae*), Spaln of D. melanogaster Proteins (purple, alignment of Ref-Seq proteins from *D. melanogaster*), RNA-Seq (alignment of Illumina RNA-Seq reads from *D. ananassae*) from adult females (red), adult males (blue), and Wolbachia-cured embryos (pink; alignment of Illumina RNA-Seq reads from *D. ananassae*), transcripts (green) including coding regions predicted by TransDecoder and splice junctions predicted by regtools using *D. ananassae* RNA-Seq (SRP006203, SRP007906, PRJNA257286, PRJNA388952). Splice junctions shown have a minimum read-depth of 17761 with >1000 supporting the reads shown. **(C) Dot plot of Glyp-PA in *D. melanogaster* (*x*-axis) vs. the orthologous polypeptide in *D. ananassae* (*y*-axis)**. Amino acid number is indicated along the left and bottom; exon number is indicated along the top and right, and exons are also highlighted with alternating colors.

### Protein Model

*Glyp* in *D. ananassae* has four coding sequences (CDSs) within the genomic sequence. The unique protein sequence (Glyp-PA; Glyp-PB) is translated from two mRNA isoforms that differ in their UTRs (*Glyp-RA*; *Glyp-RB*; Figure 1B). Relative to the ortholog in *D. melanogaster*, the CDS number and protein isoform count are conserved. The sequence of Glyp-PA in *D. ananassae* has 98.46% identity (E-value: 0.0) with the protein-coding isoform Glyp-PA in *D. melanogaster*, as determined by *blastp* (Figure 1C). Coordinates of this curated gene model are stored by NCBI at GenBank/BankIt (accession **<u>BK063018 and BK063019)</u>**. These data are also available at the TrackHub listed in “Extended Data” below.

## Methods

“Detailed methods including algorithms, database versions, and citations for the complete annotation process can be found in Rele et al. (Rele et al., 2023). Students used the GEP instance of the UCSC Genome Browser v.435 (https://gander.wustl.edu) (Kent et al., 2002; Navarro Gonzalez et al., 2021) to examine the genomic neighborhood of *Glyp* gene in the *D. melanogaster* genome assembly (Aug. 2014; BDGP Release 6 + ISO1 MT/dm6). Students then retrieved the protein sequence for the *D. melanogaster Glyp* for a given isoform and ran it using *tblastn* against the *D. ananassae* genome assembly (GCA_000005115.1) (Drosophila 12 Genomes Consortium et al., 2007) on the NCBI BLAST server (https://blast.ncbi.nlm.nih.gov/Blast.cgi, (Altschul et al., 1990) to identify potential orthologs. To validate the potential ortholog, students compared the local genomic neighborhood of the *D. ananassae* ortholog with the genomic neighborhood of *Glyp* in *D. melanogaster*. This local synteny analysis included the two upstream and downstream genes relative to the putative *D. ananassae* ortholog. They also explored other sets of genomic evidence using multiple alignment tracks in the Genome Browser, including BLAT alignments of RefSeq Genes, Spaln alignment of *D. melanogaster* proteins, and multiple gene prediction tracks (e.g., GeMoMa, Geneid, Augustus), and modENCODE RNA-Seq from the target species. Genomic structure information (e.g., CDSs, intron-CDS number and boundaries, number of isoforms) for the *D. melanogaster* reference gene was retrieved through the Gene Record Finder (https://gander.wustl.edu/~wilson/dmelgenerecord/index.html) (Gruys et al., 2025; Rele et al., 2023). Approximate splice sites within the target gene were determined using Combined Splice Junction Predictions (Kim et al., 2015) and *tblastn* using the CDSs from the *D. melanogaste*r reference gene. “Coordinates of CDSs were then refined by examining aligned modENCODE RNA-Seq data, and by applying paradigms of molecular biology such as identifying canonical splice site sequences and ensuring the maintenance of an open reading frame across hypothesized splice sites. Students then confirmed the biological validity of their target gene model using the Gene Model Checker (https://gander.wustl.edu/∼wilson/dmelgenerecord/index.html) (Rele et al., 2023), which compares the structure and translated sequence from their hypothesized target gene model against the *D. melanogaster* reference gene model. Three independent models for this gene were generated by students under mentorship of their faculty course instructors. These models were then reconciled by a third independent researcher mentored by the project leaders to produce the final model presented here. Note: comparison of 5 ‘ and 3 ‘ UTR sequence information is not included in this GEP CURE analysis.” (Gruys et al., 2025)

## Discussion

Here, we propose a gene model for the *D. ananassae* ortholog of the *D. melanogaster* Glycogen phosphorylase (*Glyp*) gene. The genomic region of the ortholog corresponds to the uncharacterized protein XP_001962117.1 (Locus ID LOC6497423) in the Agencourt dana- caf1/DanaCAF1 Genome Assembly of *D. ananassae* (GCA_000005115.1 (Graveley et al., 2011)). This model is based on RNA-Seq data from *D. ananassae* (SRP006203, SRP007906, PRJNA257286, PRJNA388952) and *Glyp* in *D. melanogaster* using FlyBase release FB2024_02 (GCA_000001215.4) (Gramates et al., 2022; Jenkins et al., 2022; Larkin et al., 2021).

## Supporting information

GlyP Gene Model Data Files

## Extended Data

File Description: “FNA” Resource Type: “Model”

File Description: “FAA” Resource Type: “Model”

File Description: “GFF” Resource Type: “Model”

Track Hub URL: https://gander.wustl.edu/cgi-bin/hgTracks?db=DanaCAF1&lastVirtModeType=default&lastVirtModeExtraState=&virtModeType=default&virtMode=0&nonVirtPosition=&position=scaffold_12916%3A6337491-6344005&hgct_customText=track%20type=bigGenePred%20visibility=pack%20bigDataUrl=http://genemodels01.ua.edu/trackhub_hosting/models/DanaCAF1/DanaCAF1.bb

## Acknowledgements

We would like to thank Laura K. Reed and Wilson Leung, who created and maintained the GEP technological infrastructure. Thank you to FlyBase for providing the definitive database for Drosophila melanogaster gene models.

